# Lymphatic vessel transit seeds precursors to cytotoxic resident memory T cells in skin draining lymph nodes

**DOI:** 10.1101/2023.08.29.555369

**Authors:** Taylor A. Heim, Austin C. Schultz, Ines Delclaux, Vanessa Cristaldi, Madeline J. Churchill, Amanda W. Lund

**Affiliations:** Ronald O. Perelman Department of Dermatology, NYU Grossman School of Medicine, New York, NY, USA; Department of Molecular Microbiology and Immunology, Oregon Health & Science University, Portland, OR; Department of Pathology, NYU Grossman School of Medicine, New York, NY, USA; Laura and Isaac Perlmutter Cancer Center, NYU Grossman School of Medicine, New York, NY, USA

**Keywords:** resident memory, lymph nodes, infection, migration, lymphatic vessels

## Abstract

Resident memory T cells (T_RM_) provide rapid, localized protection in peripheral tissues to pathogens and cancer. While T_RM_ are also found in lymph nodes (LN), how they develop during primary infection and their functional significance remains largely unknown. Here, we track the anatomical distribution of anti-viral CD8^+^ T cells as they simultaneously seed skin and LN T_RM_ using a model of skin infection with restricted antigen distribution. We find exquisite localization of LN T_RM_ to the draining LN of infected skin. LN T_RM_ formation depends on lymphatic transport and specifically egress of effector CD8^+^ T cells that appear poised for residence as early as 12 days post infection. Effector CD8^+^ T cell transit through skin is necessary and sufficient to populate LN T_RM_ in draining LNs, a process reinforced by antigen encounter in skin. Importantly, we demonstrate that LN T_RM_ are sufficient to provide protection against pathogenic rechallenge. These data support a model whereby a subset of tissue infiltrating CD8^+^ T cells egress during viral clearance, and establish regional protection in the draining lymphatic basin as a mechanism to prevent pathogen spread.

**One Sentence Summary:** T cell egress out of virally infected skin via afferent lymphatic vessels seeds CD8^+^ resident memory T cells in the draining lymph node.

## Introduction

Long term cellular immunity is achieved by multiple memory CD8^+^ T cells populations defined by their distinct proliferative, cytotoxic, and migratory capabilities^1^. Circulating memory, generally categorized into central (T_CM_) and effector (T_EM_) memory, migrate through lymph and blood, respectively, allowing them to broadly survey for systemic pathogen reencounter. In contrast, resident memory T cells (T_RM_) are absent from blood and are not thought to actively circulate between tissues^2^. T_RM_ are derived from effector T cells and are the predominant memory T cell population found in non-lymphoid tissues (NLT) after pathogen clearance^3–5^. While T_RM_ surveil relatively modest regions compared to circulating memory, they excel at cytotoxicity, can constitutively express granzyme B^6,7^, and activate local immunity through cytokine production ^8^. Therefore, the seeding of T_RM_ within specific anatomic compartments enforces local, rapid protection upon pathogen rechallenge and can be protective in the setting of cancer^9,10^. The likelihood that T_RM_ may also underlie diverse pathologies^11^, highlights an ongoing need to fully define the mechanisms that determine T_RM_ formation, maintenance, and reactivation in diverse tissues.

Interestingly, T_RM_ also form in lymph nodes (LN)^12–15^, but exactly when LN T_RM_ form following primary infection, and if there is a specific pattern of distribution has yet to be fully defined. Anatomically mapping the establishment of LN T_RM_, however, is limited by the use of systemic viral infection models, the persistence of chronic antigen, and other confounding factors such as the longevity of T_RM_ in the upstream NLT. For instance, a significant population of antigen-specific T_RM_ emerge in the lung-draining, mediastinal LN following influenza infection in mice^15^. Lung T_RM_, however, are poorly maintained^16^ and this transient nature may be what allows repositioning to the draining LN (dLN) following resolution of influenza infection in the lung^15^. Furthermore, much of what is known regarding LN T_RM_ comes from experiments where established T_RM_ in skin and mucosa are reactivated with cognate peptide, which then mobilizes these established T_RM_ to dLNs, where they form LN T_RM7,13,17_. Still, like their NLT counterparts, LN T_RM_ stand to play a role in rapid local pathogen control. Therefore, a mechanistic understanding for where and when LN T_RM_ form, the path precursor populations take, and their functional contribution relative to similarly positioned T_CM_, will be necessary to determine the impact of this regional memory population in health and disease.

To functionally and ontologically map the *de novo* formation of LN T_RM_ following primary infection, we used a model infection of skin, vaccinia virus (VV), that is both acute and local^18^. Using this model, we demonstrate that VV-specific LN T_RM_ form coincident with T_RM_ in the infected skin, are spatially restricted forming exclusively in the dLN, depend upon afferent lymphatic vessel transport, and are remarkably stable over time even in the absence of cognate antigen. Antigen presentation and priming in the dLN does not appear sufficient to generate LN T_RM_. Instead, their formation depends on the migration of CD8^+^ T cells from infected skin to dLNs via dermal lymphatic vessels. Importantly, we demonstrate that LN T_RM_ are transcriptionally poised for rapid effector function and are sufficient to mediate local viral control in the LN upon rechallenge. Our data support a model whereby contraction of peripheral tissue anti-viral responses coincides with seeding of resident memory in the LN, establishing an anatomically restricted, long-lived depot of regional immunity.

## RESULTS

### Lymph node resident memory T cells form in draining lymph nodes following skin infection

To track the formation of LN T_RM_ and their anatomical distribution we used the VV model of skin infection by scarification. When administered by scarification, VV resolves within 15 days, and viable virus stays constrained to the initial site of infection^18–20^. As a function of the restricted pattern of infection, antigen presentation and adaptive immune responses depend upon lymphatic transport to the dLN^21^. To track the formation, distribution, and contraction of antigen-specific CD8^+^ T cell responses, we adoptively transferred naïve CD90.1^+^ P14 T cells (CD8^+^ TCR transgenic T cells specific for H2-D^b^ GP_33-41_) one day prior to infection via scarification on the ear skin with VV expressing GP_33-41_ (VV-GP33). At 28 days post infection (d.p.i.), P14 T cells were present at higher numbers and frequencies in the dLN (parotid) compared to the contralateral, non-draining LN (ndLN) (**Fig 1A and B**). The enrichment for P14 T cells in dLNs, even a month after infection, indicated the possibility that a subset was resident. Consistent with residence, ∼25% of P14 T cells in the dLN expressed CD69, while P14 T cells in the ndLN expressed very little surface CD69 and CD69^+^CD62L^-^ P14 T cells were over 20-fold more prevalent in the dLN than the ndLN (**Fig 1C and D**). These CD69^+^CD62L^-^ P14 T cells bared greater phenotypic similarity to CD69 expressing cells in the skin (skin T_RM_) than T_CM_ in the same dLN. Further consistent with a resident phenotype^14^, they expressed some CD103, high levels of CXCR6, and were low for TCF1 (**Fig 1E**).

**Figure 1.**
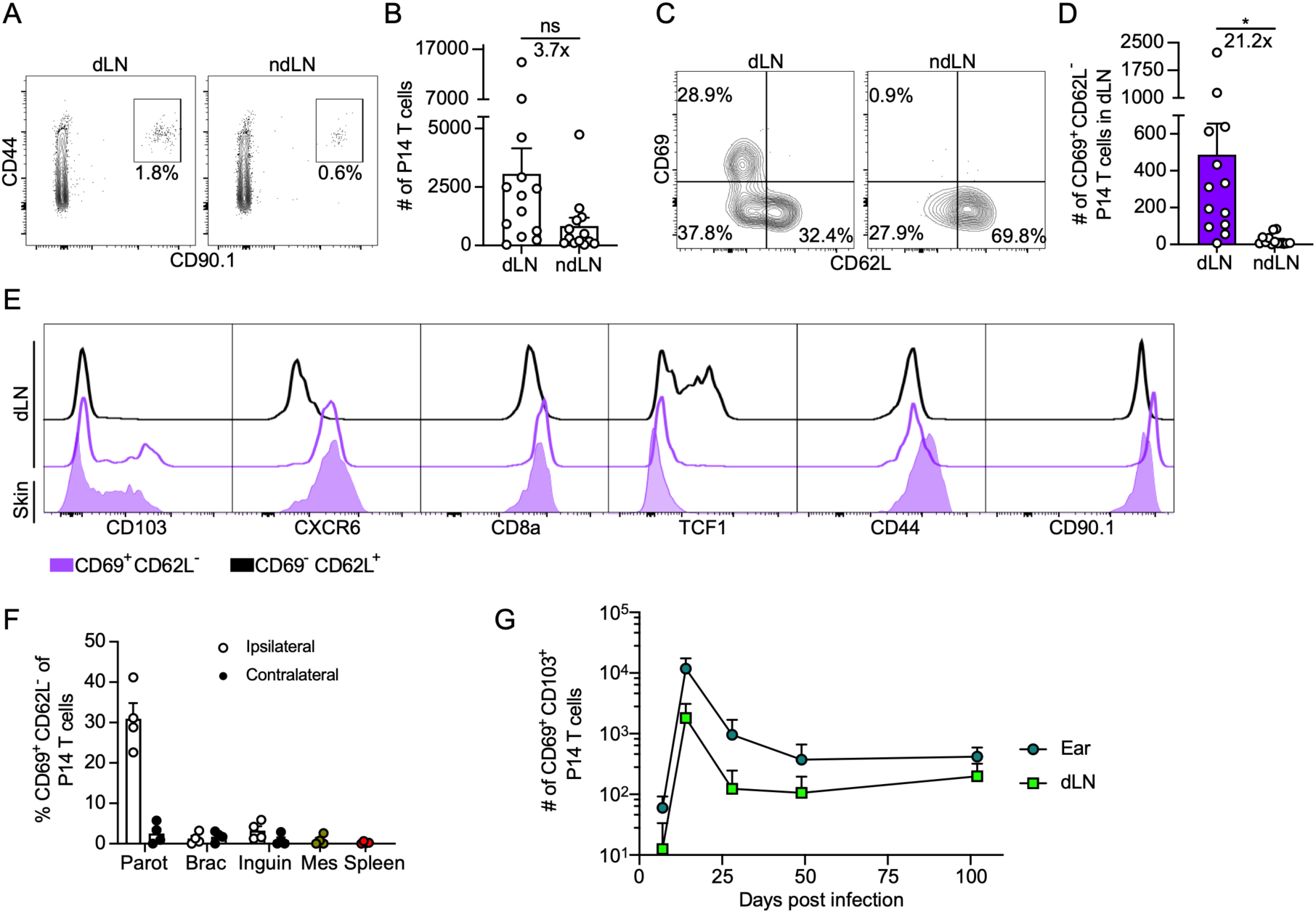
Lymph node resident memory T cells form in draining lymph nodes following skin infection. (**A-G**) P14 TCR-tg T cells (CD90.1^+^) were transferred into naïve mice (CD90.1^-^) and the following day, ear skin was infected by scarification with vaccinia virus expressing GP33_33-41_ (VV-GP33). (**A**) Representative flow plots (left, gated on live, CD8α^+^, CD45 IV^-^, lymphocytes) and quantification (**B**) of P14 T cells in lymph nodes (LN) 28 days post infection (d.p.i.). Draining LN (dLN, ipsilateral parotid); non-draining LN (ndLN, contralateral parotid). (**C**) Representative flow plots (left, gated on CD44^+^CD90.1^+^ cells in (**A**)) and quantification (**D**) of CD69^+^CD62L^-^ P14 T cells in LNs 28 d.p.i. (**E**) Representative histograms at 28 d.p.i. comparing P14 T cells. Top is CD69^-^CD62L^+^ P14 T cells in the dLN; middle and bottom are CD69^+^CD62L^-^ in dLN (middle) or skin (bottom). (**F**) Percent CD69^+^CD62L^-^ P14 T cells in ipsilateral and contralateral parotid (Parot), brachial (Brac), and inguinal (Inguin) LNs as well as mesenteric (Mes) LNs and spleen 28 d.p.i. (**G**) Number of CD69^+^CD103^+^ P14 T cells in skin and draining parotid LN over time. (**A**-**E**) Data are combined from 4 experiments for a total of n=13 mice. (**F**) Data are representative of two experiments with 4 mice per experiment. (**G**) Data are combined from at least two experiments per time point with 3-4 mice per experiment. Day 102 time point represents one experiment at day 101 and another at day 103 post infection. Bars represent average + SEM. Each point represents an individual mouse. Statistical significance determined using paired student’s t test. * p ≤ 0.05, ** p ≤ 0.01, *** p ≤ 0.001, **** p ≤ 0.0001

At memory time points, CD69^+^CD62L^-^ P14 T cells appeared uniquely in the dLN and were absent from all other secondary lymphoid organs examined, including the contralateral parotid LN, mesenteric LN and spleen (**Fig 1F**), consistent with a non-circulating, resident phenotype. To functionally test their independence from circulating populations in the blood, we administered 100 μg αCD62L to mice 26 d.p.i. (VV expressing SIINKFEL; VV-OVA), to block new lymphocyte recruitment to the LN, and 2 days later we measured the number of CD69^+^CD62L^-^ TCR-tg T cells in the dLN (CD45.1^+^ OT-1 T cells). Consistent with the hypothesis that these cells were resident, blockade of CD62L did not alter the number of LN T_RM_ in the dLN compared to IgG treated controls (**Fig S1A and B**) indicating that LN T_RM_ populations were likely not maintained via continuous recruitment from the blood. Therefore, at memory time points, CD69^+^CD62L^-^ antigen specific CD8^+^ T cells will be referred to as LN T_RM_.

We next wanted to determine the temporal kinetics of LN T_RM_ formation. To better distinguish between LN T_RM_ and CD69 expressing cells undergoing priming at early time points after infection we tracked a subset of LN T_RM_ expressing CD69 and CD103. CD69^+^CD103^+^ P14 T cells were largely absent from the dLN as well as the skin at day 7 but emerged progressively over time (**Fig 1G**). P14 T cells in the dLN expressing CD69 and CD103 were similarly maintained over time compared to P14 T cells with the same phenotype in the skin and could still be found over 100 d.p.i. (**Fig 1G**). We found that CD103^+^ P14 T cells in the dLN were not specifically localized within the dLN. Instead, CD103^+^ P14 T cells were present in most compartments of the dLN including the subcapsular sinus, paracortex and interfollicular regions, and interestingly, also the perinodal fat (**Fig S1C**). Taken together, these data indicate that localized, primary viral infection in skin seeds long-lived T_RM_ at the local site of infection and simultaneously in the immediate dLN.

### Dermal lymphatic vessels are necessary for LN T_RM_ formation after skin infection

We next asked if the initial formation of LN T_RM_ depended upon dermal lymphatic transport, which physically links skin to the dLN. K14-VEGFR3-Ig (K14-V3) mice express a soluble VEGFR3-Ig fusion protein that effectively traps VEGF-C and prevents the development of dermal lymphatic vessels in skin^22^. We transferred naïve P14 T cells into K14-V3 or WT mice and infected ear skin the following day with VV-GP33. Mice were sacrificed at least 48 d.p.i.. Interestingly, the number of skin T_RM_ (CD69^+^CD62L^-^) was similar between K14-V3 and WT mice (**Fig 2A**), indicating that despite the complete lack of lymphatic transport from skin, K14-V3 infected mice eventually initiate antigen-specific CD8^+^ T cell responses that traffic into skin, as we demonstrated previously^21^. LN T_RM_, however, were almost entirely absent from K14-V3 mice (**Fig 2B-D**), demonstrating a critical role for lymphatic transport between skin and the dLN for LN T_RM_ establishment. The altered priming and viral kinetics observed in K14-V3 mice^21^, however, may disrupt the normal development of memory. To normalize both priming of antigen-specific CD8^+^ T cells and viral kinetics, we transferred naive P14 T cells into K14-V3 or WT mice and infected with LCMV (intraperitoneal) the following day. 35 days post LCMV infection, the ear skin was challenged with VV-GP33 and mice were sacrificed at least 40 days later (**Fig 2E**). LCMV immune K14-V3 and WT mice showed similar VV-GP33 viral loads within infected ears at 6 and 11 d.p.i. (**Fig S2A**). Intriguingly, the number of skin T_RM_ was significantly higher in K14-V3 mice (**Fig 2F**), potentially indicating that lymphatic vessel mediated egress out of skin limits the magnitude of skin T_RM_. Like the primary VV-GP33 infection, however, LN T_RM_ were still completely absent from K14-V3 dLNs (**Fig 2G and H**). To ask this a third way, we normalized the primary effector response by co-infection with VV-GP33 (ear skin via scarification) and LCMV (intraperitoneal) (**Fig S2C**). We again found much fewer CD69^+^CD62L^-^ P14 T cells in the LNs of K14-V3 mice 60 d.p.i. (**Fig S2D and E**). These findings reveal that LN T_RM_ that form following cutaneous VV infection are dependent upon lymphatic transport from skin.

**Figure 2.**
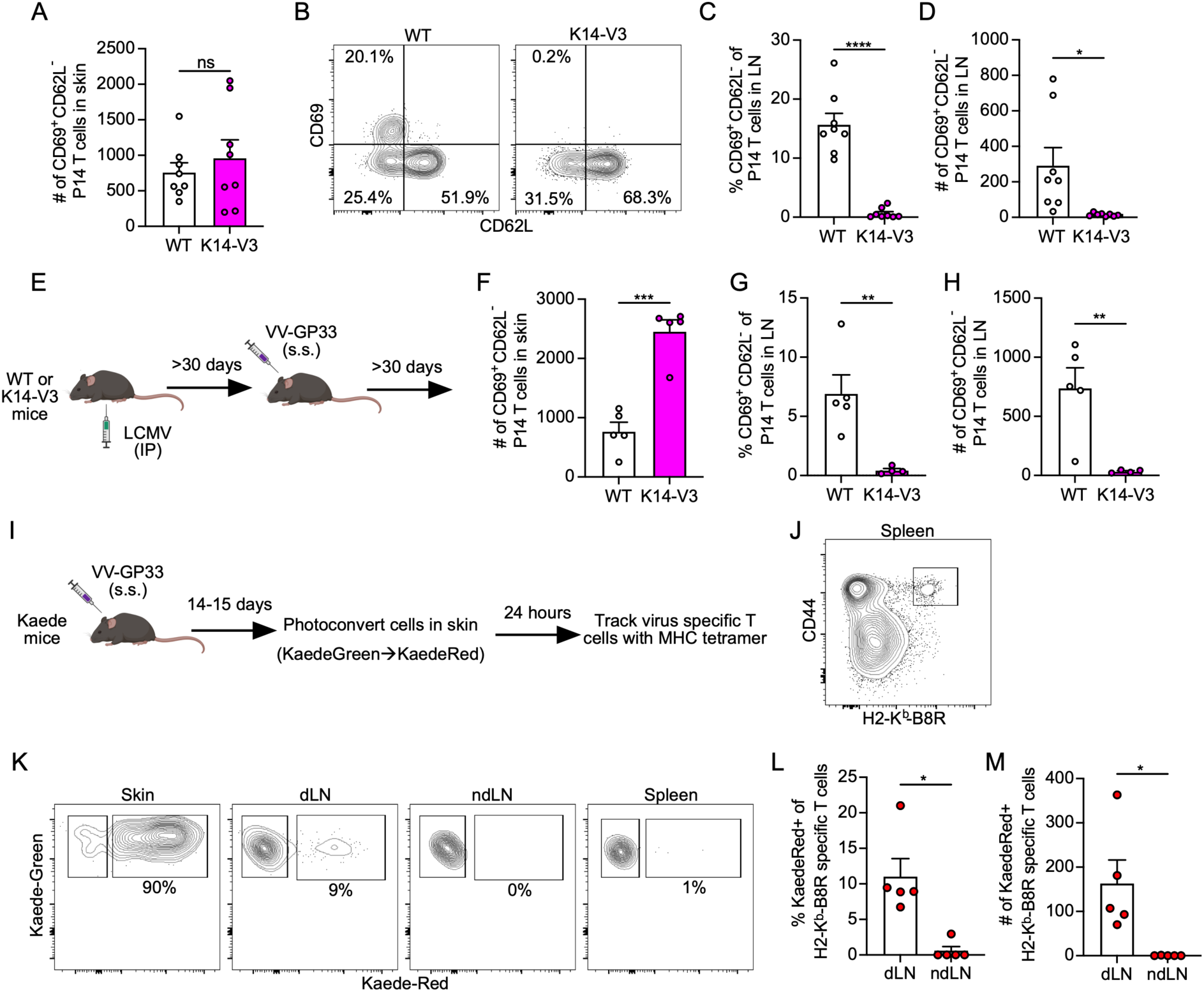
Dermal lymphatic vessels are necessary for LN TRM formation after skin infection. (**A-D**) CD90.1^+^ P14 TCR-tg T cells were transferred to WT or K14-VEGFR3-Ig (K14-V3) mice and the following day ear skin was infected with vaccinia virus expressing GP33_33-41_ (VV-GP33) via scarification. (**A**) Number of CD69^+^CD62L^-^ P14 T cells in the skin 49-50 days post infection. (**B**) Representative flow plots gated on live, CD44^+^CD8α^+^CD90.1^+^ lymphocytes and (**C**) quantification of percent and (**D**) total number of CD69^+^CD62L^-^ in draining lymph node (dLN). Data are combined from 2 experiments with 3-5 mice per group per experiment. (**E**) Experimental design for (**F**-**H**) where CD90.1^+^ P14 T cells were transferred to WT or K14-V3 mice and infected with LCMV the following day. After 30 days, mouse ears were challenged with VV-GP33 and rested at least 30 days before sacrifice. Data are representative of three experiments with 3-5 mice per group per experiment. (**F**) Number of CD45 IV^-^CD90.1^+^CD69^+^CD62L^-^ P14 T cells in skin. (**G**) Percent and (**H**) number of CD45 IV^-^CD90.1^+^CD69^+^CD62L^-^ P14 T cells in dLN. (**I**) Experimental design for (**J-M**) where photoconvertible Kaede-tg mice were infected with VV-GP33. Infected ears were photoconverted 14-15 days later and sacrificed 24 hrs later. (**J**) Representative flow plot of tetramer staining for H2-K^b^-B8R-specific T cells. Gated on live, CD8α^+^ lymphocytes. (**K**) Representative flow plots and (**L**) quantification of percent and (**M**) number of Kaede-Red fluorescence in tetramer^+^ cells. Data are representative of two experiments with 5 mice per experiment. Statistical significance determined using unpaired student’s t test (**A,C,D,F,G,H**) or paired student’s t test (L,M). Bars represent average + SEM. Each point represents an individual mouse. * p ≤ 0.05, ** p ≤ 0.01, *** p ≤ 0.001, **** p ≤ 0.0001

While lymphatic vessels are required to prime adaptive immune responses in dLNs, CD8^+^ T cells also access peripheral tissue lymphatic vessels to migrate away from sites of inflammation^13,23,24^ and peripheral tissue T_RM_ downregulate lymphatic homing mechanisms to enforce their residence^25–27^. We therefore asked if antigen-specific CD8^+^ T cells egressed from infected ear skin at effector time points. We tracked the egress of VV-GP33-specific CD8^+^ T cells using MHCI tetramer (H2-K^b^-B8R_20-27_) in Kaede transgenic mice, which express a photoconvertible fluorescent protein KaedeGreen that converts to KaedeRed upon exposure to 405nm light^13,20,23,24^. Kaede mice were infected with VV-GP33 and 15-16 days later endogenous leukocytes in skin were photoconverted (**Fig 2I** and **J**). One day after photoconversion, ∼10% of all B8R-specific CD8^+^ T cells in the dLN were KaedeRed^+^ indicating that VV-specific T cells egressed at effector time points from infected skin (**Fig 2K-M**). These data together raised the possibility that the failure to form LN T_RM_ in mice lacking dermal lymphatic vessels could be explained by the inability to reposition T_RM_ precursors from infected skin to the dLN.

### Egressing T cells share transcriptional similarities with LN T_RM_

If antigen-specific CD8^+^ T cells egressing out of skin were the precursors to LN T_RM_, we posited that they might already exhibit a transcriptional program aligned with their residence potential. Therefore, to understand the cell states that egressed infected skin at effector time points, we sorted polyclonal CD8^+^CD44^+^ KaedeRed and KaedeGreen T cells (negative for intravascular staining) from LNs following photoconversion 11 d.p.i. with VV-GP33 and performed scRNAseq. Prior to sequencing, we also tracked antigen specificity by labeling virus-specific cells with two MHCI tetramers (H2-K^b^-B8R_20-27_ and H2-D^b^-GP_33-41_) conjugated to APC. These labeled cells were then marked with an oligonucleotide tagged antibody against APC via cellular indexing of transcriptomes and epitopes (CITE-Seq) allowing resolution of a subset of the antigen-specific repertoire within the polyclonal population. Transcriptional analysis and visualization by Uniform Manifold Approximation Projection (UMAP) projection of the integrated data (KaedeGreen and KaedeRed) identified 5 clusters (**Fig 3A**). Our CITE-Seq approach identified the majority of tetramer^+^ cells to be in clusters 0, 4 and 3 (**Fig 3B**). Egressing CD8^+^ T cells (KaedeRed) were most highly enriched in clusters 0 and 4, while KaedeGreen, LN populations were enriched in clusters 1 and 2 but represented in all clusters (**Fig 3C and D**). The KaedeGreen clusters represented central memory-like (cluster 1: *Cd44*, *Tcf7*, *Sell*), naïve (cluster 2: *Tcf7*, *Ccr7*, low *Cd44*), and proliferating (cluster 3: *Mki67* and *Top2a*) CD8^+^ T cell populations. In contrast, the egressing, antigen-specific clusters (0 and 4) were defined by high expression of effector molecules (*Gzmb, Gzmk, Ifng*) and transcripts associated with resident memory (*Cxcr6, Itga1, Id2*) (**Fig 3E and F**). Interestingly, despite their effector phenotype and expression of transcripts associated with residence (e.g. *Cxcr6*, *Bhleh40, Id2, Itga1*), egressing cells also maintained expression of transcripts associated with circulation including *Klf2* and *S1pr1*, consistent with their recent migration from skin (**Fig 3E**). Furthermore, only a small percentage of cells in clusters 0 and 4 expressed *Itgae,* and *Cd69* was lowly expressed relative to the naïve and T_CM_ clusters (**Fig 3E**). Still, when scored for a published gene signature for LN T_RM_ ^14^the KaedeRed clusters (0 and 4) scored highest (**Fig 3G**). While these data indicated that egressing CD8^+^ T cells were unlikely to be fully differentiated T_RM_, they supported the hypothesis that egressing, effector CD8^+^ T cells could be poised for residence and thereby represent a potential precursor population that seeds T_RM_ in dLNs over time.

**Fig. 3.**
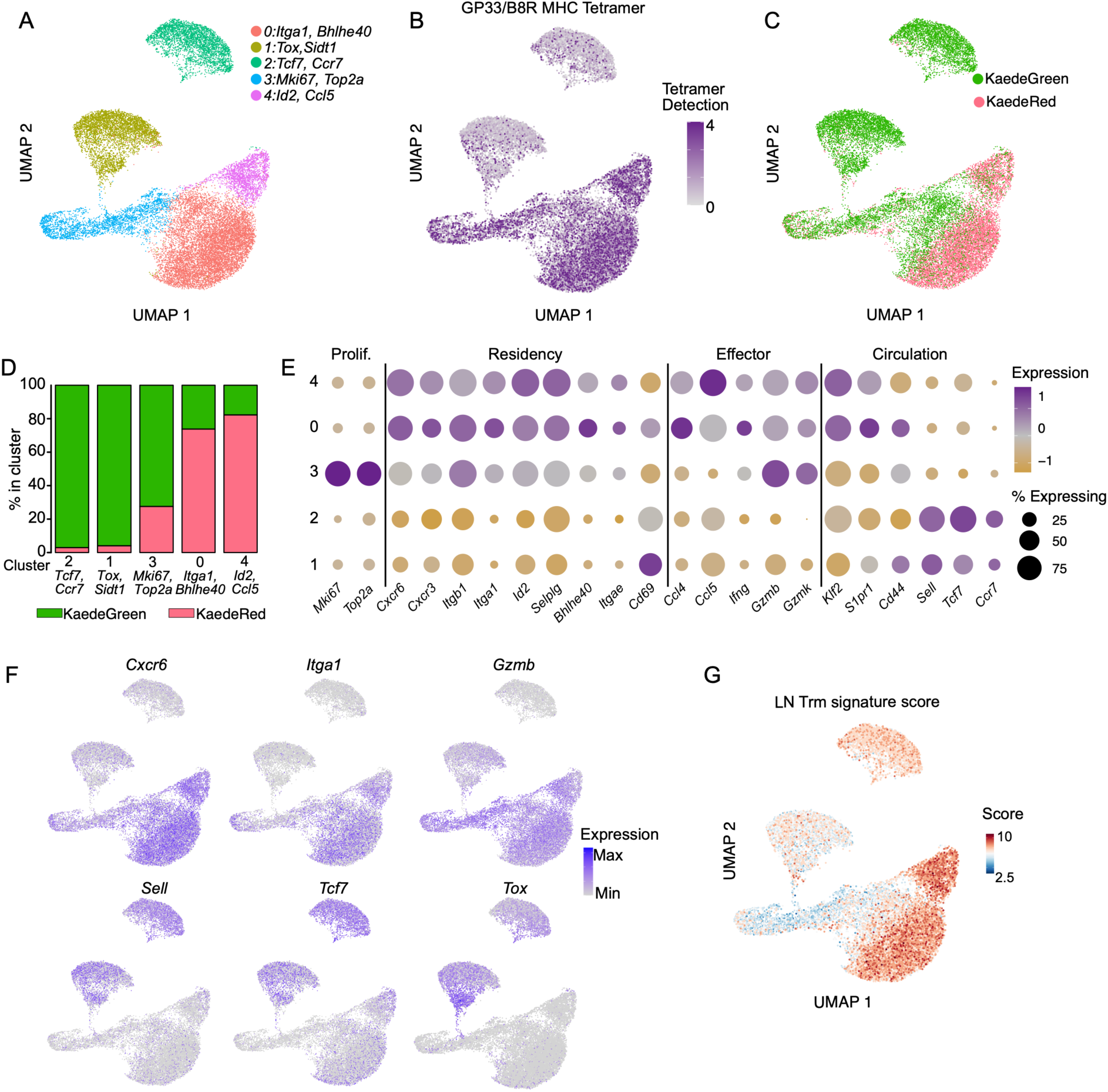
Egressing T cells share transcriptional similarities with LN T_RM_. Kaede-tg ears were infected with VV-GP33 by scarification and photoconverted (KaedeGreen → KaedeRed) 11 days later. 24 hours after photoconversion, KaedeGreen^+^ and KaedeRed^+^ cells were sorted by FACS (Live, CD45 IV^-^ CD8^+^CD44^+^) from draining lymph nodes pooled from 29 mice and analyzed by scRNA-seq. Prior to sorting, viral specific cells were stained with 2 APC conjugated tetramers (H2-K^b^-B8R_20-27_ and H2-D^b^-GP_33-41_). APC was then tagged with an oligonucleotide to identify a subset of viral-specific cells by scRNA-Seq. (**A**) UMAP projection of cells colored according to cluster. (**B**) CITE-seq detection of tetramers visualized by UMAP projection. (**C**) UMAP projection showing distribution of KaedeGreen^+^ or KaedeRed^+^ and (**D**) quantification of their proportions within each cluster. (**E**) Expression of select genes across clusters normalized from −1 to 1. Size of dot corresponds to percentage of cells in cluster expressing the transcript. (**F**) UMAP projection of select transcripts or (**G**) signature score for LN resident memory T cells (Molodtsov et al. 2021).

### T cell transit through skin is required for the formation of LN T_RM_

To directly test the hypothesis that these egressing effector T cells were LN T_RM_ precursors, we depleted P14 T cells from the blood, and secondary lymphoid organs (1-2μg αCD90.1; **Fig 4A-C**) but left them intact in skin (**Fig S3A**)^15,28,29^ to see if effector T cells in skin were sufficient to populate LN T_RM_. Circulating T cells were still depleted 15 days later (**Fig S3B**) and there was no difference in the total number of skin T_RM_ compared to control treated mice (**Fig S3C**). As expected, however, P14 T cells in skin of mice 15 days post depletion were enriched for a T_RM_ phenotype, expressing CD69 and CXCR6 (**Fig S3D and E**). Consistent with our hypothesis, we found that P14 LN T_RM_ still formed specifically in the dLN, indicating that their precursors must have been in the infected skin at time of depletion (**Fig. 4D**). These P14 T cells in the dLN 15 days post depletion were also enriched for a LN T_RM_ phenotype (CD69, CXCR6, CD103) compared to control treated mice (**Fig 4E and F**). Importantly, when all circulating CD8^+^ T cells were depleted 10 days following VV-GP33 infection, we saw a similar rebounding of endogenous LN T_RM_ specific to the dLN as tracked by tetramer (**Fig S3F-H**).

**Figure 4.**
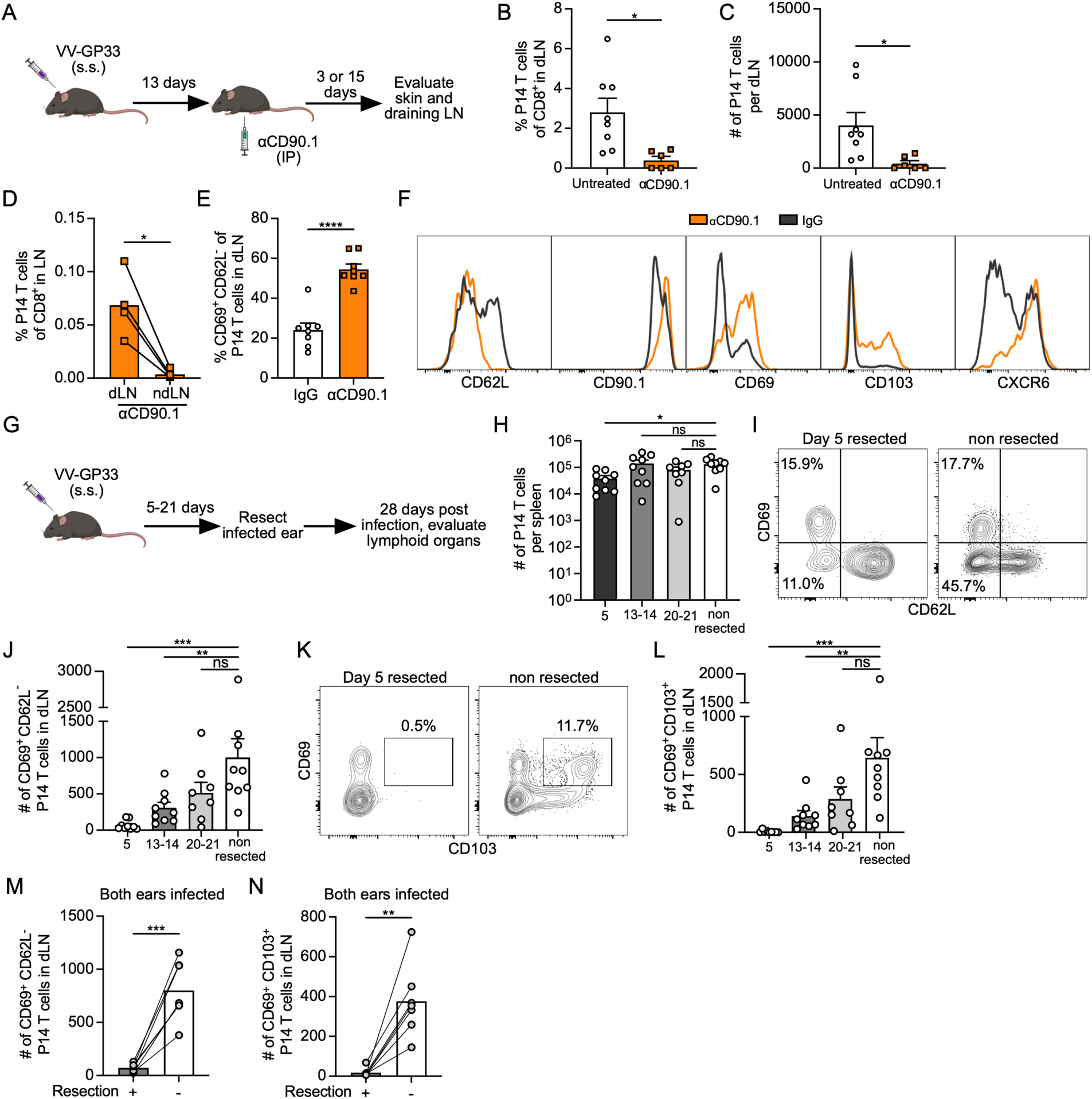
T cell transit through skin is required for the formation of LN T_RM_. (**A**) Experimental design for (**B**-**F**) where P14 TCR-tg T cells (CD90.1^+^) were transferred into naïve mice (CD90.1^-^) and the following day ear skin was infected by scarification with vaccinia virus expressing GP_33-41_ (VV-GP33). At 13 days post infection (d.p.i.), mice were treated with IgG or αCD90.1 to deplete P14 T cells in blood and secondary lymphoid organs and sacrificed 3 or 15 days later. (**B**) Frequency and (**C**) number of all P14 T cells in untreated (14 d.p.i.) or depleted mice (16 d.p.i., 3 days post depletion) in draining lymph node (dLN). (**D**) Prevalence of P14 T cells in LNs of depleted mice 15 days post depletion (contralateral, non-draining LN, ndLN). (**E**) Percent of P14 T cells with a CD69^+^CD62L^-^ residence phenotype and (**F**) representative surface phenotype of P14 T cells in dLNs of depleted (orange) or IgG treated (black). Data are combined from two experiments (**B**,**C**,**E**,**F**) or representative of two experiments (**D**) with 3-4 mice per group per experiment. (**G**) Experimental design for (**H**-**L**) where P14 T cells were transferred into naïve mice and the following day ear skin was infected with VV-GP33. At variable times, the infected ear was resected and mice were all sacrificed 28 d.p.i. Data are combined from two experiments with 3-5 mice per group per experiment. (**H**) Total number of P14 T cells per spleen 28 d.p.i. (**I**) Representative flow plots and (**J**) quantification of CD69^+^CD62L^-^ P14 T cells in dLNs. (**K**) Representative flow plots and (**L**) quantification of CD69^+^CD103^+^ P14 T cells in dLNs. For M and N, P14 T cells were transferred into naïve mice and the following day skin from both ears were infected with VV-GP33. One ear was resected at 5 d.p.i. and dLNs from the resected and unresected ear were compared 28 d.p.i.. Quantification of CD69^+^CD62L^-^ (**M**) or CD69^+^CD103^+^ (**N**) P14 T cells in dLNs. Data are combined from two experiments (**M,N**) with 3-4 mice per group per experiment. Bars represent average + SEM. Statistical significance was determined using unpaired student’s t test (**B**,**C**,**E**), paired student’s t test (**D,M,N**) or one way ANOVA (**H**,**J**,**L**). * p ≤ 0.05, ** p ≤ 0.01, *** p ≤ 0.001, **** p ≤ 0.0001

These data indicated that at least a subset of LN T_RM_ formed from T cells that had migrated out of infected skin, but it did not exclude the possibility that some LN T_RM_ were seeded directly at priming. To ask whether priming was sufficient to seed LN T_RM_, we resected infected ear skin at 5,13 or 20 d.p.i. and quantified T cell expansion and LN T_RM_ formation at 28 d.p.i. (**Fig 4G**). Equivalent levels of circulating P14 T cell memory formed by day 28 across all groups relative to the unresected with the exception of the day 5 resected group where we observed a significant but only modest decrease in total P14 T cells (**Fig 4H**). With this model, we were thus able to investigate LN T_RM_ formation in mice where P14 T cells had undergone similar priming and expansion in the dLN but were unable to reposition from the skin at effector time points. We found that the total number of CD69^+^CD62L^-^ P14 T cells in dLNs were significantly reduced with ear resection (**Fig 4I-J**) and only increased the longer the ear skin was left intact. Interestingly, there was an even more severe loss of CD69^+^CD103^+^ P14 T cells in the dLN that also increased with time (**Fig 4K and L**). To further validate these results, we repeated the day 5 resections in a co-infection setting where both ears were simultaneously infected, but only a single ear was resected. Again, LN T_RM_ failed to form in the LN draining the resected tissue site, despite normalized systemic memory and inflammation, while T_RM_ formed in the contralateral dLN (**Fig 4M and N**). These data indicated that LN T_RM_ abundance correlated with available time for T cell transit from skin to the dLN. These experiments altogether, therefore, support the conclusion that T cell egress is both necessary and sufficient for LN T_RM_ formation post VV-infection.

### LN T_RM_ establishment is dependent upon antigen recognition in skin

Recognition of antigen by CD8^+^ T cells augments T_RM_ formation in skin after VV infection ^18^. However, the role of antigen recognition and where antigen recognition might take place, has yet to be fully understood for LN T_RM_. Our previous data indicated that primary antigen encounter in the dLN was not sufficient for LN T_RM_ formation. Therefore, to test if antigen recognition in skin by CD8^+^ T cells boosts LN T_RM_, we infected mice on opposite ears with VV or VV-OVA the day after CD45.1^+^ OT-1 T cell transfer (**Fig 5A**). At 28 d.p.i. OT-1 T cells were present at higher rates (% CD45.1^+^ of CD8^+^) in the VV-OVA dLN compared to the VV dLN in the same mouse (**Fig 5B and C**). Furthermore, OT-1 T cells in the VV-OVA dLN were more likely to be CD69^+^CD62L^-^ than OT-1 T cells in the VV dLN (**Fig 5D and E**), indicating that a second antigen encounter in either the VV-OVA infected skin or dLN promoted LN T_RM_ formation.

**Figure 5.**
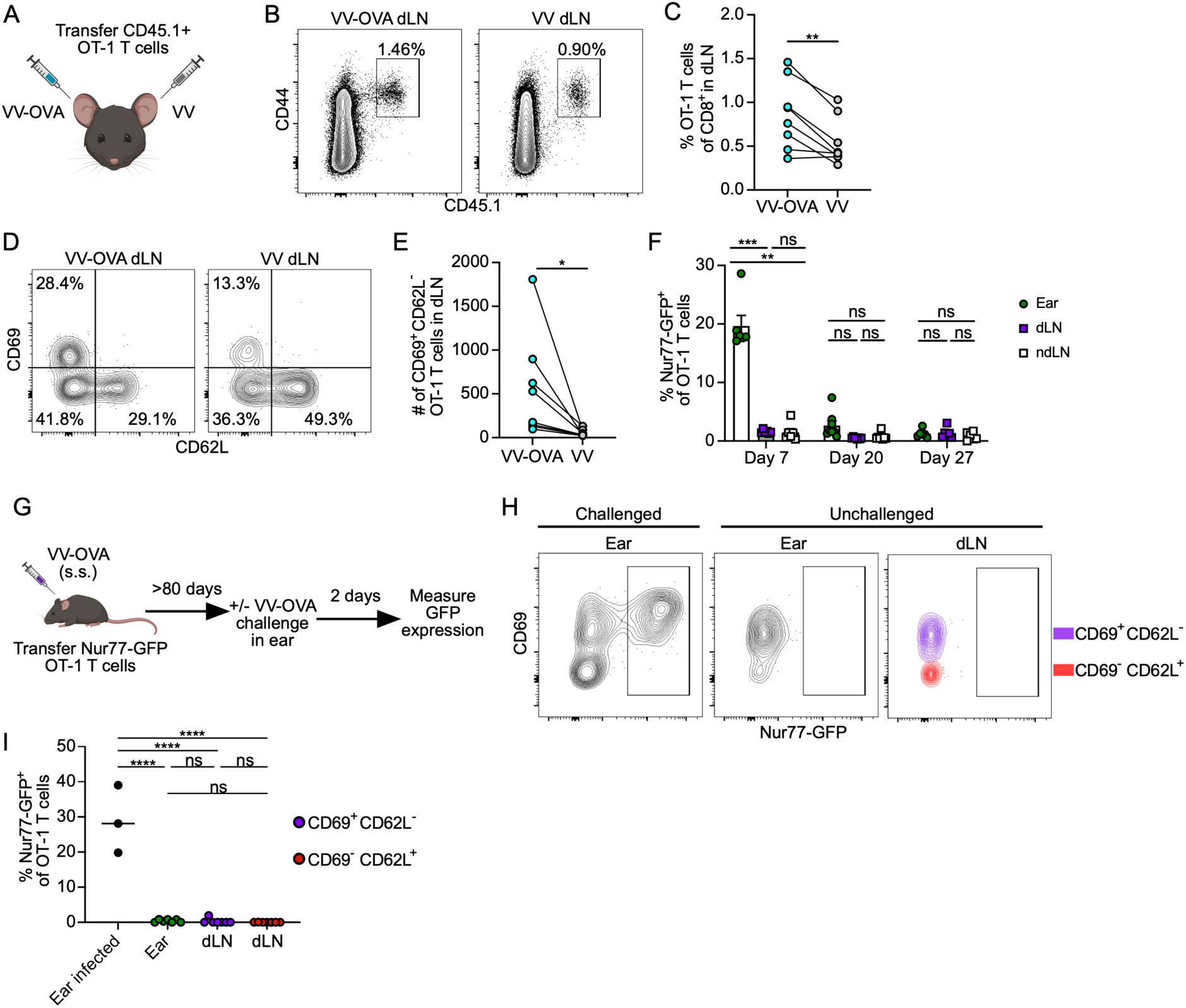
LN T_RM_ establishment is dependent upon antigen recognition in skin. (**A**) Experimental design for (**B**-**E**) where CD45.1^+^ OT-1 TCR-tg T cells were transferred to naïve mice. The next day mice were infected on the right ear skin with vaccinia virus (VV) and left ear with VV expressing SIINFEKL (VV-OVA) via scarification and sacrificed 27-28 days later. (**B**) Representative flow plots and (**C**) quantification of frequency of CD45.1^+^ OT-1 T cells 27-28 d.p.i. in draining lymph nodes (dLN). Flow plots are gated on live, CD8α_+_ lymphocytes. (**D**) Representative flow plots gated on CD45.1+ OT-1 T cells in dLN and (**E**) quantification of number of CD69+CD62L-OT-T T cells in dLNs. Data are combined from two experiments with 4 mice per experiment. (**F**) Quantification of GFP expression in CD45.1^+^ OT-1 Nur77-GFP T cells transferred to mice one day prior to infection with VV-OVA. Skin, dLN, and non-draining LN (ndLN). (**G**) Experimental design for (**H** and **I**) where CD45.1^+^ OT-1 Nur77-GFP T cells were transferred into naïve mice and the ear was infected the following day with VV-OVA. 80 d.p.i mice either received a secondary VV-OVA challenge (Challenged) or did not (Unchallenged). (**H**) Representative flow plots and (**I**) quantification of GFP expression in CD45.1^+^ OT-I T cells. Data are combined from 2 experiments. Each point represents an individual mouse. Statistical significance determined using paired student’s t test (**C**,**E**) or one way ANOVA (**H**). * p ≤ 0.05, ** p ≤ 0.01, *** p ≤ 0.001, **** p ≤ 0.0001

Based on the dependence of skin transit and antigen encounter for the formation of LN T_RM_, we posited that antigen encounter could occur in skin prior to tissue egress. Our kinetic data indicated that the first LN T_RM_ emerge between day 7 to day 13. To track antigen encounters within this window of time, we used T cells expressing a fluorescent reporter of TCR ligation to map the probability of antigen encounter in skin and dLN. In mice with transgenic expression of green fluorescent protein (GFP) under control of the Nr4a1(Nur77) promoter (Nurr77-GFP), GFP expression faithfully and transiently reports TCR ligation^8,30^. We transferred CD45.1^+^ Nurr77-GFP OT-1 T cells into naïve mice and infected one ear the following day with VV-OVA. We measured GFP expression at 7, 20 and 27 d.p.i.. As anticipated, expression of GFP was highest in the ear at 7 d.p.i. when viral loads are still high in the skin. Interestingly, and consistent with our ear resection experiments that argue for a very early window for priming, GFP expression in the dLN was consistently low and comparable to the ndLN at all examined time points (**Fig 5F**). Anti-viral T cells in the dLN, therefore, were not actively engaging cognate peptide at any time point after 7 days. Consistent with their memory phenotype, LN T_RM_ at late memory time points (86-87 d.p.i.) remained GFP negative, indicating persistence independent of continuous antigen stimulation, similar to circulating T_CM_ (**Fig 5G-I**). Skin re-challenged with VV-OVA provides a positive control for antigen recognition (**Fig 5G**). In summary, our data is consistent with a model whereby antigen recognition during the effector phase in skin acts as a necessary second hit that boosts LN T_RM_ levels in the dLNs.

### LN T_RM_ are poised for cytotoxicity and provide localized protection

Our data highlights the simultaneous presence of antigen-specific circulating and resident memory T cell populations in the dLN after VV-GP33 infection. Whether there’s specific value to maintaining both populations, however, is unclear. To better understand how LN T_RM_ functionally differed from circulating memory populations in the dLN, we performed single cell RNA sequencing on all transferred P14 T cells in the dLN 45 d.p.i. with VV-GP33. Transcriptional analysis and visualization by UMAP projection identified 6 clusters, including a small, proliferating cluster of cells (cluster 4) defined by high transcriptional levels of cell cycling genes *Mki67* and *Top2a* (**Fig 6A**). To identify the putative T_RM_ cluster, we scored our cells with an established gene signature for LN T_RM_ in murine models of melanoma^14^. Cluster 2, characterized by high expression of *Itgae* (CD103), *Itga1* (CD49a), *Cxcr6*, and *Id2* exclusively scored for the resident memory transcriptional program (**Fig 6B**). In contrast, clusters 0, 1 and 3 possessed transcriptional programs consistent with circulating memory T cells and were defined by elevated *S1pr1*, and *Sell* (**Fig S4A-C**). Relative to the circulating clusters, cluster 2 expressed low levels of *S1pr1*, *Sell*, and *Klf2,* genes frequently associated with circulation, and scored highly for TGFβ signaling^31^ (**Fig S4A-C**), which promotes the transition to residence^32^. To further validate the predicted states of our clusters, we investigated the transcriptional regulatory networks using single-cell regulatory network inference and clustering (SCENIC) (**Fig 6B**). Interestingly, Cluster 2 scored highly for the Bhlhe40 regulon, a transcription factor recently demonstrated to be critical for mitochondrial fitness of T_RM_ (**Fig 6B**)^33^. These transcriptional changes were consistent with the identification of Cluster 2 as bona fide LN T_RM_.

**Figure 6.**
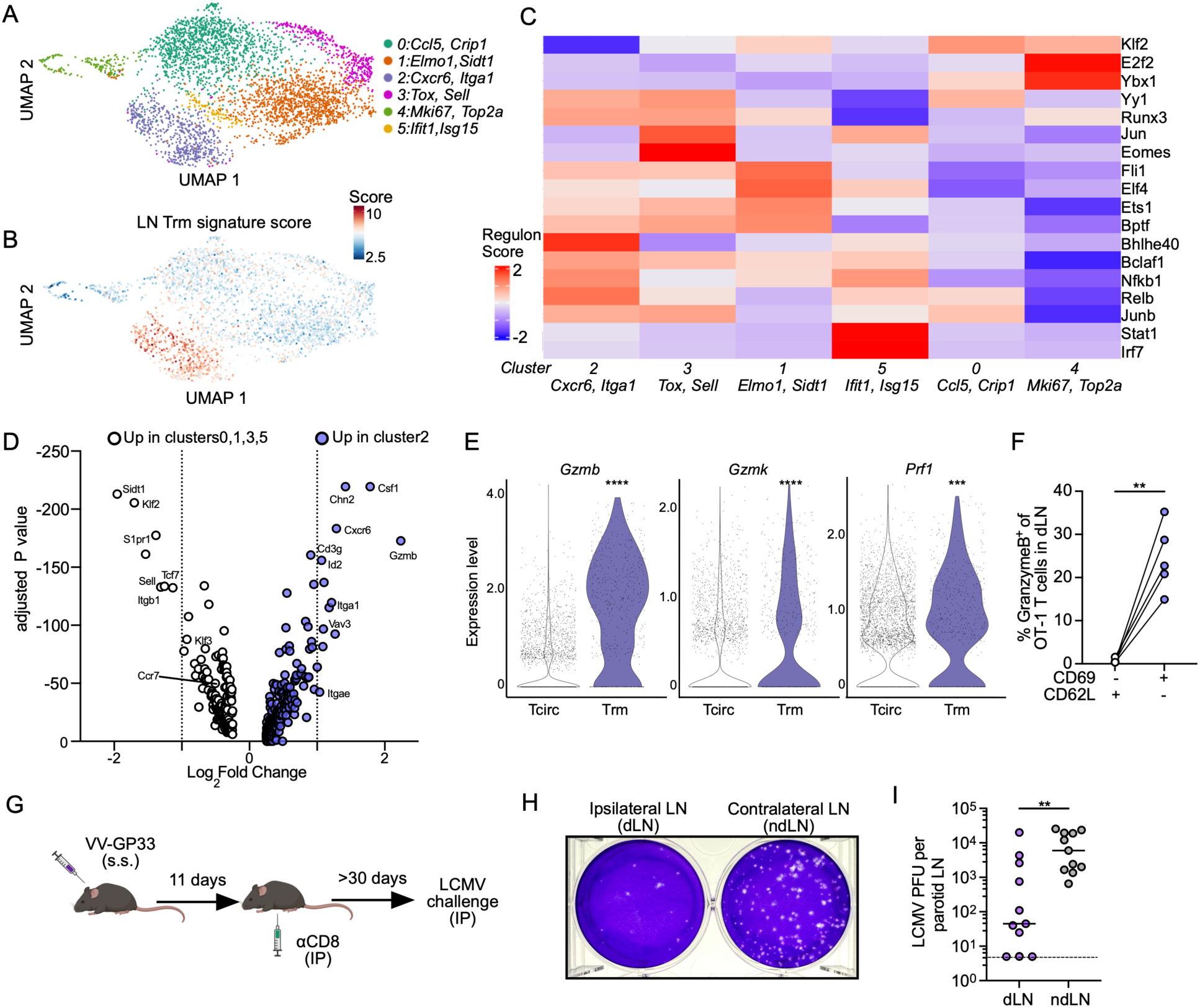
LN T_RM_ are poised for cytotoxicity and localized protection. (**A**) UMAP representation of single-cell RNA-seq data from CD45 IV^-^ P14 TCR-tg T cells isolated from draining lymph nodes (dLN) 45 d.p.i. with VV expressing GP33_33-41_ (VV-GP33). (**B**) Cells scored against a published gene signature for LN resident memory T cells (T_RM_). (**C**) Single-cell regulatory network inference and clustering (SCENIC) analysis inferring gene regulatory networks in each cluster. (**D**) Differentially expressed genes between resident (cluster 2) and circulating memory (clusters 0,1,3,5) cells. (**E**) Expression of *Gzmb*, *Gzmk*, and *Prf1* in T_CIRC_ (clusters0,1,3,5) and T_RM_ (cluster 2). (**F**) Expression of GranzymeB by OT-1 TCR-tg T cells at least 60 days after ear skin infection with VV expressing SIINFEKL (VV-OVA) as measured by flow cytometry. Data are representative of two experiments with 5 mice per experiment. (**G**) Experimental design for (**H** and **I**) where naïve mice were infected on one ear with VV-GP33 and treated with 100μg of αCD8α 11 days after infection. At least 30 days later mice were challenged with lymphocytic choriomeningitis virus (LCMV) via intraperitoneal injection. (**H**) Representative image of plaques and (**I**) quantification of plaque forming units (PFU) measured 3 days following challenge dLN (ipsilateral) or contralateral, non-draining LN (ndLN). Data are combined from 3 experiments with 3-4 mice per experiment. Each point represents an individual mouse. Statistical significance determined using pairwise Wilcoxon rank test (**D**,**E**) or paired student’s t test (**F**,**I**). ** p ≤ 0.01, *** p ≤ 0.001, **** p ≤ 0.0001

We then asked whether additional differences in expression might indicate altered functional potential. Differential gene expression analysis revealed that transcripts of genes associated with TCR signaling and cytotoxicity tended to be more highly expressed in LN T_RM_ (cluster 2) than other circulating clusters (T_CIRC_; clusters 0,1,3 and 5) (**Fig S4D**). *Gzmb*, *Gzmk*, and *Prf1* transcript levels were all higher on the LN T_RM_ cluster than T_CIRC_ clusters (**Fig 6E**). We validated elevated levels of GranzymeB by flow cytometry on OT-1 LN T_RM_ (CD69^+^CD62L^-^) relative to OT-1 T_CM_ (CD69^-^CD62L^+^) from the same dLN at >60 days after VV-OVA infection in the ear skin (**Fig 6F**). These findings indicated that LN T_RM_ were specifically poised for cytotoxicity compared to circulating memory CD8^+^ T cells in the same LN. Intriguingly, OT-1 T_RM_ in the dLN and skin expressed comparable protein levels of GranzymeB (**Fig S4E**), which is consistent with the high degree of phenotypic similarities between skin T_RM_ and LN T_RM_ and further supports the model of a common progenitor or differentiation program between these populations.

To directly test whether the LN T_RM_ were sufficient to provide local protection against viral rechallenge in LNs, we infected mice with VV-GP33 on a single ear, and then depleted circulating cytotoxic T cells with anti-CD8α antibody 11 d.p.i. (**Fig S3E-G**). As we demonstrated previously (**Fig. S3E-G**), this experimental setup establishes endogenous anti-viral LN T_RM_ in the absence of circulating memory. At least 30 days after depletion, mice were rechallenged with systemic LCMV via intraperitoneal injection and LCMV viral loads were measured in the dLN and contralateral ndLN 3 days after challenge (**Fig 6G**). We found that LNs with T_RM_ (ipsilateral parotid to initial VV-GP33 infection) had ∼100 fold lower viral titers relative to contralateral, ndLNs in the same mouse (**Fig 6H and I**). Taken together, these data indicate that LN T_RM_ are sufficient to provide rapid protection to safeguard LNs from re-infection.

## Discussion

LNs are essential for initiating *de novo* adaptive immune responses to infections and maintaining homeostasis through multiple mechanisms of peripheral tolerance. While many studies have investigated CD8^+^ T cell immunosurveillance of non-lymphoid tissues, CD8^+^ T cell immunosurveillance of LNs is poorly understood. Here we detail the ontogeny and developmental cues needed for the formation of a resident CD8^+^ T cell population capable of rapid pathogen control in the LN. We find that the formation of LN T_RM_ depends upon lymphatic transport and specifically the trafficking of effector CD8^+^ T cells from peripheral infected tissue that happens simultaneous with resident T cell differentiation in skin. By tracking local antigen recognition, we find that LN T_RM_ formation depends on a secondary antigen encounter in skin. Upon egress to LNs, these newly resident memory cells are uniquely poised for cytotoxicity, thereby providing swift and localized protection of secondary lymphoid organs after systemic viral challenge.

Optimal adaptive immune responses to pathogens depend on presentation of pathogen-derived antigens in the LN, a process mediated by afferent lymphatic vessels^21^. Afferent lymphatic vessels transport lymph containing antigens, antigen presenting cells, and in some cases viable pathogens into the subcapsular sinus of the LN^34–36^. Pathogen dissemination to the LN, however, has the potential to impair ongoing or future adaptive immune responses^37^. Peripheral tissue lymphatic vessels can themselves provide an active barrier to viral dissemination via lymph^20,38^, and if this barrier is breached, subcapsular sinus macrophages scavenge lymph-borne pathogenic material to prevent further systemic dissemination^34^. The presence of these overlapping mechanisms to restrict pathogen infiltration into the LN further supports the concept that protecting the LN from direct infection may help to optimize adaptive immunity and limit systemic spread. While immunosurveillance of LNs has been thought to be performed by circulating memory T cells, T_RM,_ which provide rapid protective immunity in NLTs, are also found in both murine and human LNs. Indeed, in HIV-infected humans, CD8^+^ T cells resembling T_RM_ are present in lymphoid tissues where they correlate with improved viral control^39^. Furthermore, high expression of a transcriptional signature associated with LN T_RM_ in metastatic LNs correlates with improved survival in humans with metastatic melanoma^14^. Here we demonstrate that LN T_RM_ form in the draining LNs following primary infection and rapidly act to control the local infiltration of virus upon rechallenge. This work therefore adds to an emerging paradigm that the establishment of LN T_RM_ within lymphatic draining basins may provide important regional protection from invading pathogens and tumor cells.

While LN T_RM_ in both mouse and human LNs share many transcriptional similarities with NLT T_RM 4,40_, discerning their ontogeny is complicated by the multiple points of entry for lymphocytes into LNs (blood and lymph), and the diverse microbial experiences of adult humans. The use of systemic viral infections^13,28,39^ or autoimmune-melanoma models^14^ where antigen is systemic or chronic, respectively, further complicates the interpretation of the developmental history of LN T_RM_. Using localized and acute VV infection in mouse skin, a model with well-defined spatiotemporal kinetics of antigen encounter^18^, we demonstrate that precursors of VV-induced LN T_RM_ are recruited from infected skin via afferent lymphatic vessels specifically back to the dLN simultaneous with the formation of T_RM_ in skin. K14-V3 mice lacking dermal lymphatic vessels showed a complete inability to form LN T_RM_ after VV-GP33 infection in the skin. This was particularly striking in our co-infection models, where LCMV infects the parotid LN^41^ and still no LN T_RM_ formed in the absence of dermal lymphatic transport. Furthermore, resection of infected skin resulted in reduction of LN T_RM_ but similar numbers of VV-GP33 specific memory T cells in the spleen and LN T_RM_ still formed after depletion of VV-GP33 specific T cells at effector time points from blood and secondary lymphoid organs. These data all together support a model where LN T_RM_ are not formed during initial priming in the dLN, but rather derive specifically from precursors that emigrate from infected skin. As a result, local lymphatic transport contributes to the anatomical distribution of LN T_RM_ precursors to the first draining LN synchronous with the formation of memory in skin, which establishes a second layer of protection from subsequent rechallenge.

The signals that determine the probability of T cell egress and retention remain incompletely understood. However, antigen recognition by effector CD8^+^ T cells in NLT seems to enhance T_RM_ formation^18^ and directly regulates expression of surface receptors implicated in migration^23,42^. By simultaneously infecting opposing ears with VV and VV-OVA, we found that CD8^+^ T cells were more likely to differentiate into LN T_RM_ in the LN draining the ear infected with matched VV-expressing cognate antigen. Given our trafficking data, which indicated that cells must transit through infected skin to seed LN T_RM_, we hypothesized that a second antigen encounter (outside of priming) was required for LN T_RM_ formation. It remained unclear if this additional antigen encounter occurred in the infected skin or within the dLN following egress out of the skin. Reporters for TCR ligation, showed very little signs of CD8^+^ T cell recognition of antigen in the dLN at day 7 post infection and onward, arguing against dLN antigen recognition as the key signal endorsing LN residency. Rather the last point for robust antigen encounter was in skin at day 7, indicating that antigen encounter in skin may initiate the residence program that establishes upon entry to the LN. Egressing T cells showed signs of high effector capability and early transcriptional similarities to LN T_RM_, but maintained their migratory potential. Given that antigen encounter in skin might simultaneously define both skin and LN resident, what sorts effector CD8^+^ T cell between these two compartments remains unclear, but may depend on the context of the antigen encounter, who presents and where.

Interestingly, both in VV^42^ and tumor models^23^ the strength of TCR signaling regulates the expression of chemokine receptors that may define the migratory properties of effector CD8^+^ T cells. High affinity antigen encounter in skin enforces expression of CXCR6 and downregulates S1PR1 with no effect on CCR7^42,43^. While our data are consistent with the idea that antigen encounter in skin directs a subset of egressing T cells towards a LN T_RM_ fate, the specific machinery needed for their migration from skin to the dLN remains unclear. Potential candidates include receptors for sphingosine 1-phosphate (S1P), such as S1PR1 and S1PR5, and chemokine receptors, such as CCR7, and CXCR4. CCR7 has been implicated in promoting CD4^+^ T cell egress^44,45^ out of acutely but not chronically inflamed skin^46^, and CXCR4 promotes CD8^+^ T cell egress from tumors^23^. However, neither recent emigrants nor established LN T_RM_ expressed high levels of *Ccr7 or Cxcr4*. S1P gradients are thought to direct T cell egress out of NLTs, based in large part on the high expression of CD69, a S1PR1 antagonist, in established T_RM_. Consistent with a role for S1P gradients in residence, forced expression of S1PR1 reduces NLT T_RM 47_, while loss of S1PR5 leads to reduced numbers of NLT T_RM 26_. The impact of CD69 loss on residence is less clear, with some evidence for decreased CD8^+^ T cell accumulation in skin^32^, but no impact in many other NLTs except for kidney^48^. Here, we found that recent emigrants, even those scoring for a T_RM_-like signature, expressed *S1pr1* while, as expected, established LN T_RM_ did not. This may support the idea that S1P gradients guide CD8^+^ T cell exit at effector time points, but whether S1PR1 is specifically required or not needs to be formally tested. Interestingly, while both recent emigrants from inflamed skin and established LN T_RM_ express high levels of *Cxcr6*, CXCR6 is dispensable for both CD8^+^ T cell entry into and egress out of VV infected skin, and instead supports interstitial survival^43^. Further work, therefore, is required to deconvolve the tissue and context-specific chemotactic machinery necessary for CD8^+^ T cell egress from infected skin.

In summary, here we define the developmental and migratory pathway governing T_RM_ formation in skin-draining LNs during primary infection. We demonstrate that CD8^+^ T cells egressing from inflamed skin are the precursors to LN T_RM_, and therefore that lymphatic transport is critical to seed a potent, cytotoxic population capable of restricting systemic dissemination of pathogens in dLNs. As such, we propose that lymphatic transport compartmentalizes the resident memory repertoire creating anatomical layers of protective immunity to restrict pathogen spread and systemic disease.

## Acknowledgements

We thank Katherine S. Ventre for technical assistance and members of the Lund lab for helpful discussion. AWL is supported by the National Institutes of Health (R01CA238163), the Cancer Research Institute (Lloyd J. Old STAR Award), the American Cancer Society (RSG-18-169-01-LIB), and the American Association for Cancer Research (AACR-BMS Midcareer Female Investigator Grant). TAH is supported by the National Institutes of Health (T32-AI100853).

## Author Contributions

Author contributions: T.A. Heim conceived, performed, and analyzed experiments and wrote the manuscript. A.C. Schultz, I. Delclaux, and M. J. Churchill performed experiments. V.Cristaldi and A.C. Schultz performed computational analyses. A.W. Lund acquired funding; conceived, and analyzed experiments; and wrote the manuscript. All authors contributed to revision and approved final submission.

## Materials and Methods

### Mice

C57BL/6J (Strain:000664), B6.SJL-PtprcaPepcb/BoyJ (CD45.1, Strain:002014), C57BL/6-Tg(TcraTcrb)1100Mjb/J (OT-1, Strain:003831), B6.PL-Thy1a/CyJ (CD90.1, Strain:000406), C57BL/6-Tg(Nr4a1-EGFP/cre)820Khog/J (Nur77-GFP, Strain:016617) mice were purchased from Jackson Laboratory and bred in specific pathogen-free conditions at Oregon Health and Sciences University or New York University Grossman School of Medicine. P14 TCR-tg mice ^49^ were maintained in our colony and were a gift from Jeffrey Nolz. B6.Cg-Tg(CAG-tdKaede)-15Utr (Kaede-Tg) ^50^ were obtained via D.J. Fowell in agreement with RIKEN BioResource Research Center. K14-VEGFR3-Ig mice ^22^ were a generous gift from Dr. Kari Alitalo and obtained from Dr. Melody A. Swartz. Mice were age and sex matched and female mice were used unless otherwise stated. For adoptive transfer experiments, 20,000-40,000 CD8^+^ T cells from P14 or OT-1 mice were adoptively transferred into naïve female mice. For experiments with K14-VEGFR3-Ig mice, both male and female recipients were used. Mice were used between 8-16 weeks of age. All animal procedures were approved and performed in accordance with the Institutional Animal Care and Use Committees at OHSU and NYU Langone Health.

### Pathogens, infections, and plaque assays

For vaccinia virus infections (VV) by scarification, mice were infected with either VV, VV-GP33 or VV-OVA. Briefly, 2-5 x 10^6^ PFU of virus in 10μl of PBS was placed on ear pinnae and ears were poked 40-50 times with a 29-G needle. Alternatively, mice were infected with 1-2 x 10^6^ PFU of LCMV Armstrong in 200μl of PBS by intraperitoneal (I.P.) injection. Quantification of viral load was determined using standard plaque assays on Vero or BSC40 cells. Briefly, skin and LNs were mechanically homogenized in 1ml of RPMI 1%FBS and frozen at −80°C. For VV titer quantification, samples underwent 3 freeze-thaw rounds. Samples were diluted and plated on cells in 6 well plates before addition of agarose overlay. Cells were incubated for 3-4 days. Agarose was removed and plaques were visualized with crystal violet.

### In vivo antibody treatment

To deplete circulating CD90.1^+^ cells^15,28,29^ 1-2µg of αThy1.1 (HIS51) antibody in 200µl of PBS was injected I.P.. In order to deplete circulating CD8^+^ cells^51^ 100µg of αCD8α (2.43) antibody in 200µl of PBS was injected I.P.. Depletion in blood was confirmed 1 day later by flow cytometry. For CD62L blockade, mice were treated with 100µg of αCD62L (MEL-14) at 26 d.p.i. and euthanized 2 days later.

### In vivo T cell egress

To label Kaede expressing T cells, previously infected ear skin was photoconverted using 405-nm light for 150 seconds (75 seconds each side) at 10mW^52^. Mice were euthanized the following day.

### Ear resection

Mouse ear skin was removed at the base of the ear pinna with surgical scissors. Mice were treated subcutaneously with buprenorphine (0.1mg/kg) after surgery every 12 hours for a total of 48 hours.

### Immunofluorescent microscopy

LNs were removed from mice and frozen in optimum cutting temperature freezing medium. Samples were frozen on a 2-methylbutane bath over dry ice. Frozen samples were then cut on a Leica cryostat at 7-10µm per section and fixed in acetone for 10-15 minutes. Slides were blocked for 10 minutes with 1%BSA in PBS and stained with antibodies in 1%BSA in PBS for 1 hour at room temperature. Antibodies were obtained from Biolegend and Tonbo and included B220 (RA3-6B2), CD103 (2E7), LYVE1 (ALY7), CD90.1 (OX-7). Coverslips were attached with Prolong Diamond Anti-Fade and imaged on a Keyence BX-X810 or Zeiss Axio Observer 7.

### Leukocyte Isolation

Dorsal and ventral sides of ears were separated and incubated for 30 minutes at 37 C in 1ml of HBSS (Hyclone) containing CaCl_2_ and MgCl_2_ supplemented with 125 U/ml of collagenase D (Invitrogen) and 60 U/ml of Dnase-I (Sigma-Aldrich). LNs, spleens or digested skin were smashed on a scored plate and put through a 70µm cell strainer to generate single cell suspensions. Spleens were resuspended for 2 minutes in 2ml of ammonium-chloride-potassium lysis buffer.

### Flow Cytometry

For intravenous (IV) labeling, mice were injected intravenous with 2 μg of anti-CD45 or anti-CD8 antibody 3 minutes before euthanizing mice. Single-cell suspensions were stained with antibodies in FACS buffer (1%BSA in PBS) with brilliant staining buffer for 30 minutes at 4C. Antibodies were purchased from Biolegend, Cell Signaling Technologies, Invitrogen, BD Biosciences, and Tonbo and included, CD8 (53-6.7), CD62L (MEL-14), CD45.1 (A20), CD90.1 (OX-7), CD90.2 (30-H12), CXCR6 (SA051D1), CD69 (H1.2F3), CD103 (2E7), CD44 (IM7), CD45 (30-F11), and

TCF1 (C63D9). Ghost dye 780 (ZombieRed) was used to detect dying cells. For tetramer staining, single cell suspensions were incubated with H2-K^b^-B8R_20-27_ and H2-D^b^-GP_33-41_ tetramers for 45 minutes at room temperature in the dark. Tetramers were provided by the NIH Tetramer Core. Staining of intracellular TCF1 was performed with True-Nuclear Transcription Buffer set as per manufacturer suggestions. Samples were analyzed with a BD FACSymphony and Flowjo software.

### Sorting and single-cell RNA sequencing

For data in Figure3: Ears of Kaede mice were infected with VV-GP33 and photoconverted at 11 d.p.i.. Lymphocytes were isolated from dLNs after intravenous injection of anti-CD45 antibody. CD8^+^ cells were MACS enriched. Cells were stained with two tetramers (H2-K^b^-B8R_20-27_ and H2-D^b^-GP_33-41_) conjugated to APC. Cells were then incubated with TotalSeq anti-APC (Biolegend, 408009). KaedeGreen and KaedeRed cells were sorted (CD8^+^CD44^+^CD45 IV^-^) on a BD FACS Aria II. DAPI was added prior to cell sorting to select live cells.

For data in Fig 6 and Supplementary Fig 4: Ears of C57Bl/6J mice were infected with VV-GP33 the day after receiving 20,000 P14 T cells. At 45 d.p.i., lymphocytes were isolated from dLNs after intravenous injection of anti-CD45 antibody as previously described. CD90.1^+^ cells were MACS enriched and P14 T cells (CD8^+^CD90.1^+^CD44^+^CD45 IV^-^) were sorted on a BD FACS Aria II. DAPI was added prior to cell sorting to select live cells. Samples were pooled from 7 mice.

The sorted cellular suspensions were loaded on a 10x Genomics Chromium instrument to generate single-cell gel beads in 25 emulsions. Libraries were prepared using Single cell 3’Reagent kits v3.1 (Chromium Next GEM Single Cell 3’ GEM, Library & Gel Bead Kit v3.1, 16 rxns PN-1000121;10x Genomics) and were sequenced using Illumina Novaseq 6000. Quality control was performed on sequenced cells to calculate the number of genes, UMIs and the proportion of mitochondrial genes for each cell. Cells with low number of covered genes and high mitochondrial counts were filtered out. A general statistical test was performed to calculate gene dispersion, base mean and cell coverage to use to build a gene model for performing Principal Component Analysis (PCA). Data was analyzed and visualized using Seurat packages in R. For CITE-Seq^54^ analysis, data was normalized using the CLR data normalization method is Seurat. To predict the active gene regulatory networks (transcription factors with target genes) we ran the computational pipeline SCENIC^53^ on our scRNAseq data, using as input the R object generated using Seurat. We generated a complex heatmap that shows the average regulon activity in each Seurat cluster.

### Statistics

Statistical analysis was performed using GraphPad Prism 9 software. Comparisons between two groups were conducted using paired or unpaired students t tests. Comparisons between more than two groups were conducted using one way ANOVA. Proper sample size was based off of prior experience. Statistical analysis of sequencing data was performed with pairwise Wilcoxon rank test and Bonferroni correction in R. P values are reported as follows: * p ≤ 0.05, ** p ≤ 0.01, *** p ≤ 0.001, **** p ≤ 0.0001. Error bars show the mean ±SEM.

### Figure Design

Figures with experimental designs were created using Biorender.com

### Data Availability

scRNAseq data is deposited in GEO and will be made publicly available upon acceptance (GSE241820, GSE236283) and is available upon request.

---

**Supplemental Figure 1.**
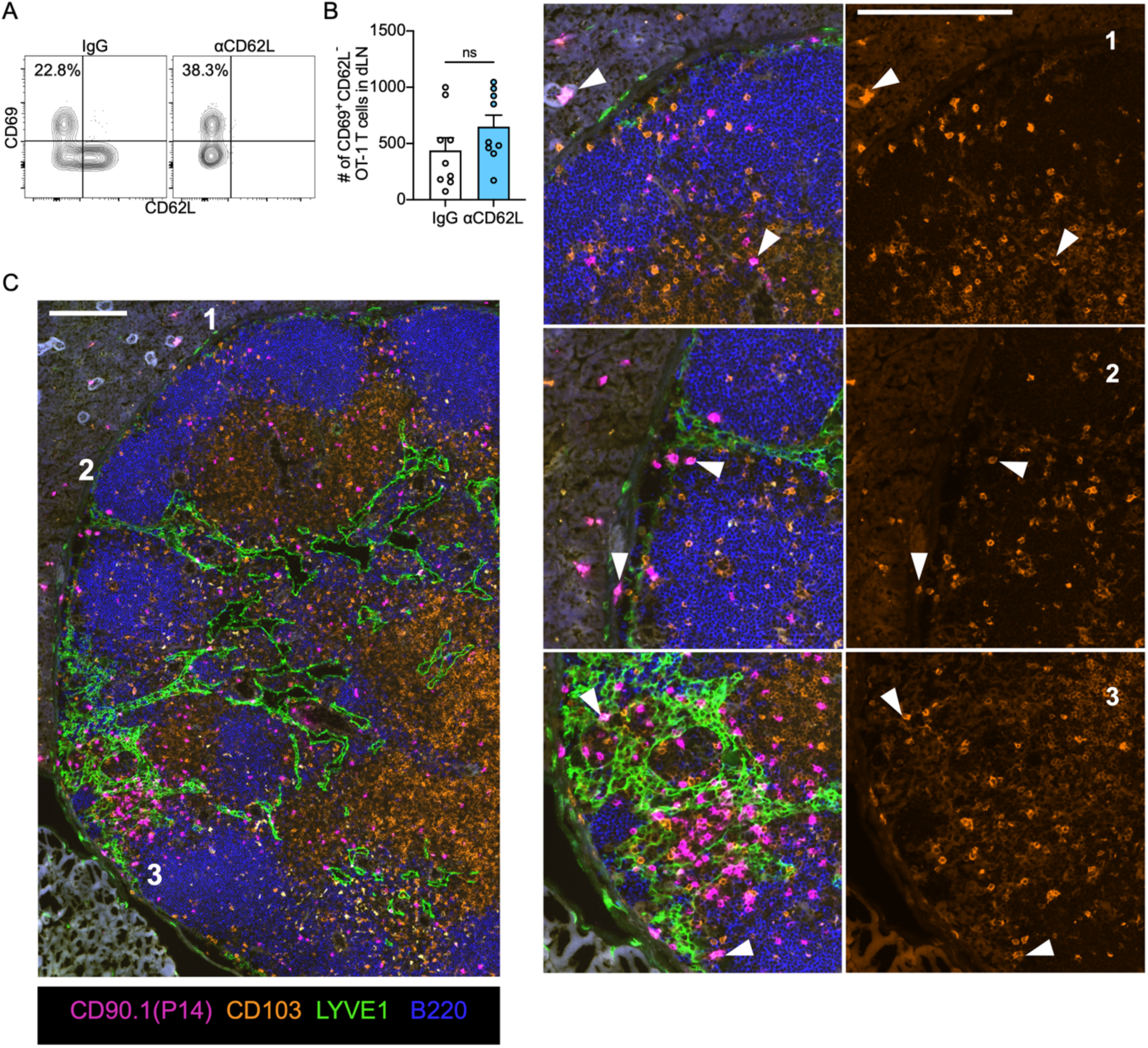
CD103^+^ P14 T cells are distributed throughout the draining lymph nodes at memory time points. CD45.1^+^, OT-1 TCR-tg T cells were transferred to naïve mice. The next day mice were infected with vaccinia virus expressing SIINFEKL (VV-OVA) via scarification. 26 days post infection (d.p.i.) mice were treated with 100 µg α-CD62L blocking antibody or IgG control and sacrificed 2 days later. (**A**) Representative flow plots of OT-1 T cells in draining lymph node (dLN) and (**B**) quantification of CD69^+^CD62L^-^ OT-I T cells in dLN (**B**). Data are combined from two experiments with 4-5 mice per group per experiment. (**C**) Representative immunofluorescent image of the dLN 47 d.p.i. with VV expressing GP33_33-41_. Insets: (1) Perinodal fat (top right) and paracortex (bottom right). (2) Subcapsular sinus. (3) Interfollicular region. Arrows denote CD90.1^+^P14 TCR-tg T cells expressing CD103. Scale bar is 200 µm. Statistical significance determined using unpaired student’s t test. Related to Figure 1

**Supplemental Figure 2.**
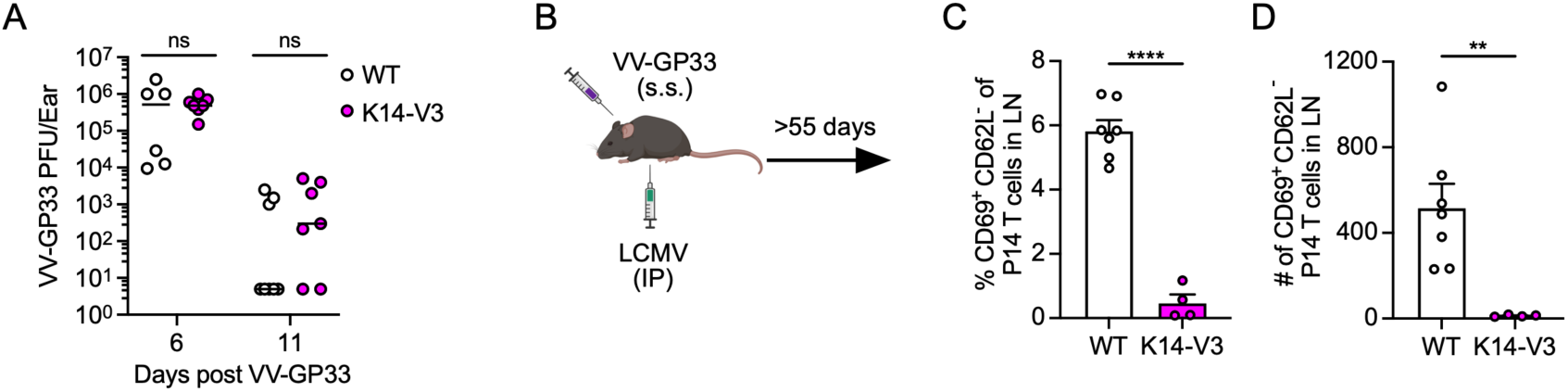
T cell egress out of skin is necessary for LN T_RM_ after vaccinia infection. (**A**) WT or K14-VEGFR3-Ig (K14-V3) mice were infected with LCMV Armstrong and challenged with VV-GP33 via scarification at least 30 days post infection (d.p.i.) Viral loads in skin were measured by plaque assay. (**B**) Experimental design for (**C** and **D**) where WT or K14-VEGFR3-Ig (K14-V3) mice were simultaneously infected with LCMV and VV-GP33 after adoptive transfer of naïve P14 T cells. (**C**) Quantification of percent and (**D**) number of CD69^+^CD62L^-^ P14 T cells >55 d.p.i.. Data are combined from 2 experiments with 2-5 mice per group per experiment. Each point represents an individual mouse. Statistical significance determined using unpaired student’s t test. ** p ≤ 0.01, **** p ≤ 0.0001 Related to Figure 2

**Supplementary Figure 3.**
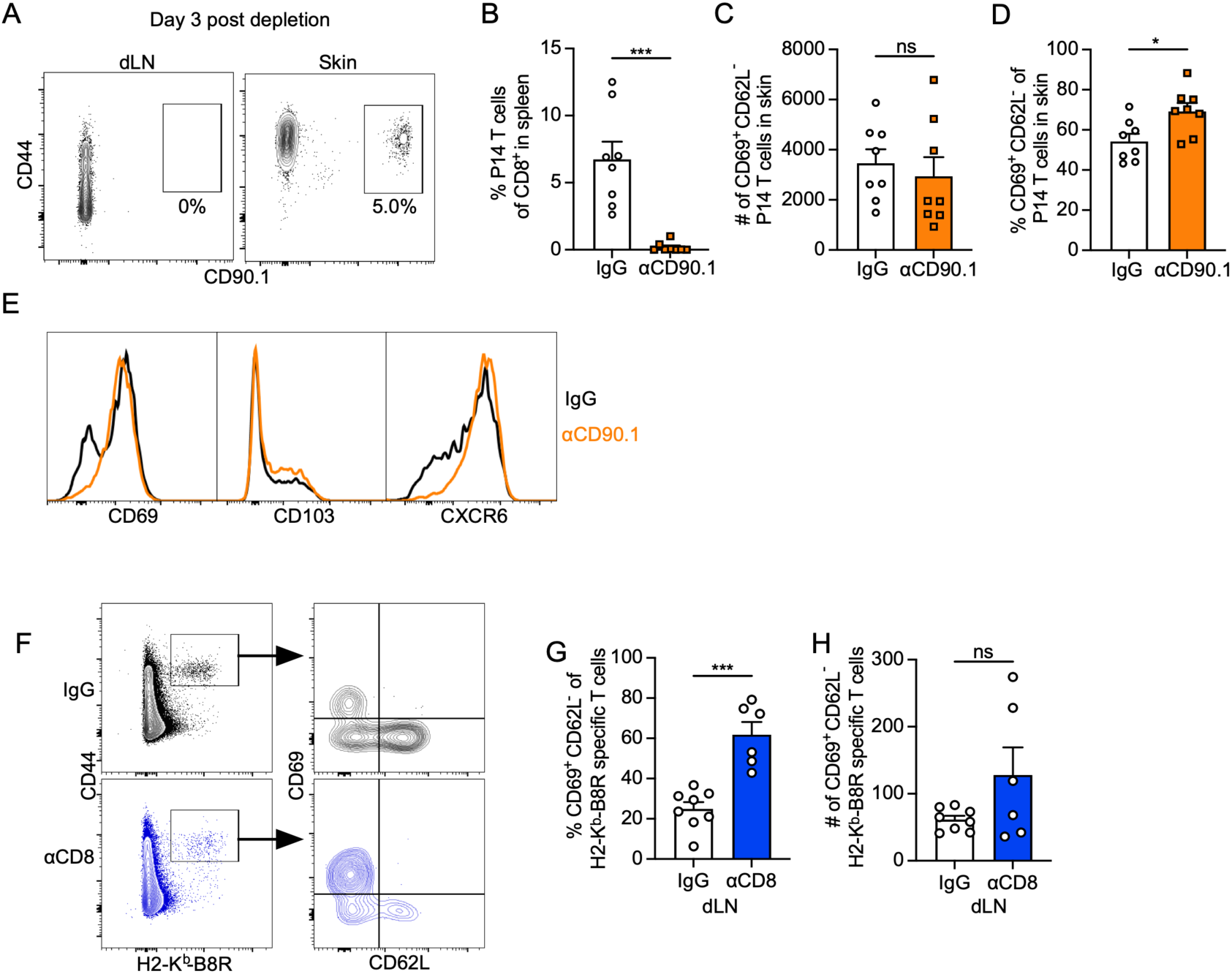
Depletion eliminates CD69^-^ P14 T cells in spleen and skin and enriches for CD69^+^ cells in skin after infection. (**A-E**) P14 TCR-tg T cells (CD90.1^+^) were transferred into naïve mice (CD90.1^-^) and the following day ear skin was infected by scarification with vaccinia virus expressing GP_33-41_ (VV-GP33). At 13 days post infection (d.p.i.), mice were treated with IgG or αCD90.1 to deplete P14 T cells in blood and secondary lymphoid organs. (**A**) Representative flow plots 3 days after treatment showing depletion of CD90.1^+^ P14 T cells in dLN but not skin. Gated on CD8^+^ lymphocytes. (**B**) Percent P14 T cells of CD8^+^ lymphocytes in spleen 15 d.p.i.. (**C**) Number and (**D**) percent of CD69^+^CD62L^-^ P14 T cells in skin 15 days post depletion. (**E**) Representative surface phenotype of P14 T cells in skin 15 days post depletion. Data are combined from two experiments with 4 mice per group per experiment. (**F-H**) Left ears of mice were infected with VV-GP33 via scarification and treated with IgG (top, black) or αCD8 (bottom, blue) at 10 d.p.i. and sacrificed at least 30 days later. (**E**) Representative flow plots of H2-K^b^-B8R tetramer staining and CD69 and CD62L expression, gated on CD8^+^ lymphocytes in the dLN. (**G**) Percent and (**H**) number of tetramer^+^CD69^+^CD62L^-^CD8^+^ lymphocytes in dLN >20 days post treatment. Data are combined from two experiments with 2-4 mice per group per experiment. Each point represents an individual mouse. Statistical significance determined using unpaired student’s t test (**A**-**C**,**F**,**G**). * p ≤ 0.05, *** p ≤ 0.001 Related to Figure 3

**Supplementary Figure 4.**
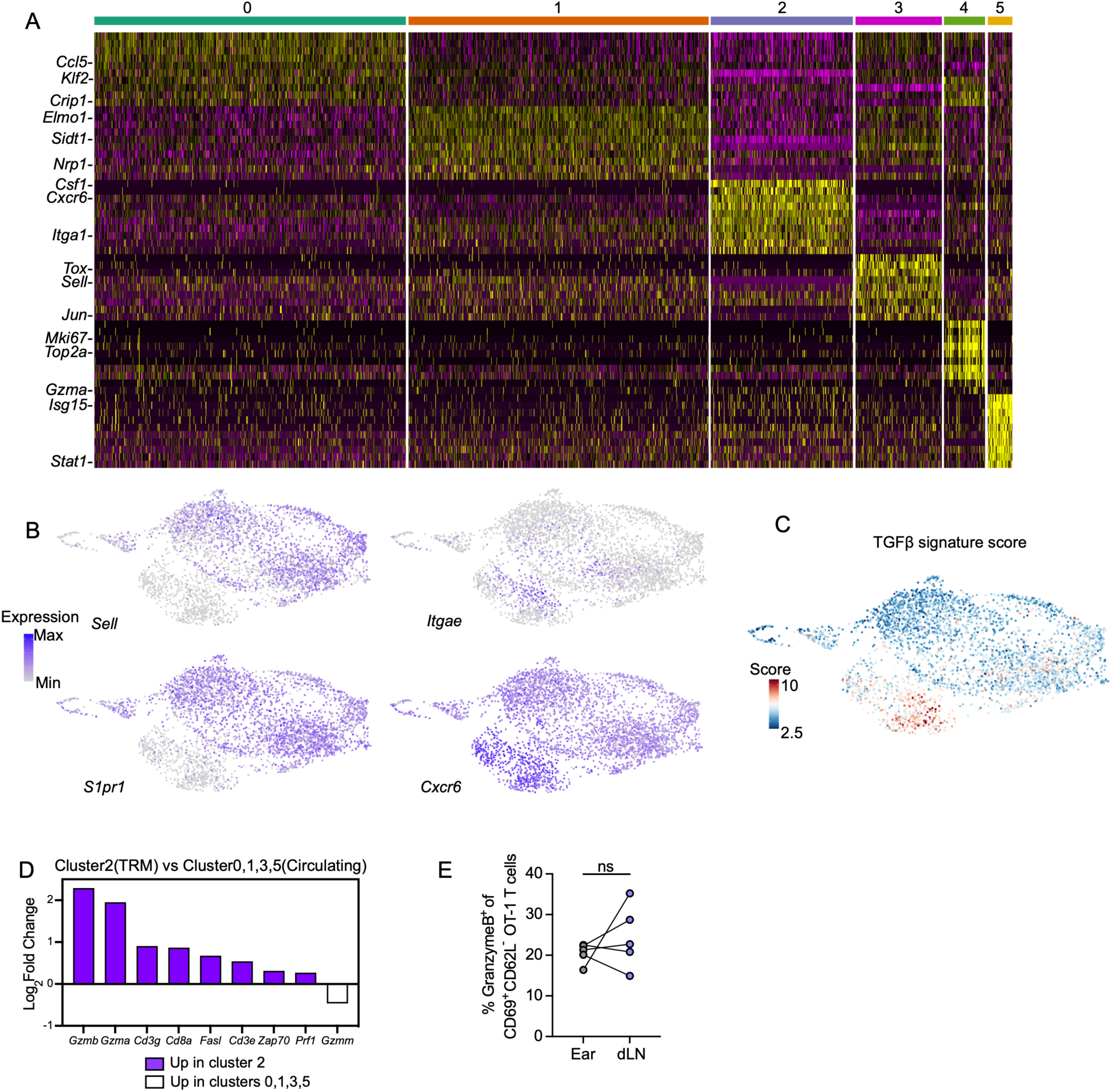
LN T_RM_ transcriptional program and effector function. (**A**) Heatmap showing the top 10 genes for each scRNAseq cluster of P14 TCR-tg T cells 45 days post infection with vaccinia virus expressing GP_33-41_ (VV-GP33) by scarification. (**B**) Feature map expression of *Sell*, *Itgae*, *S1pr1*, and *Cxcr6*. Cells were scored (**C**) for a TGFβ signaling gene signature (Nath et al 2019). (**D**) Cytotoxic transcript levels in cluster 2 relative to 0,1,3,5. (**E**) Expression of GranzymeB (GZMB) by OT-1 TCR-tg T cells at least 60 days after ear skin infection with VV expressing SIINFEKL (VV-OVA) as measured by flow cytometry. Data are representative of two experiments with 5 mice per experiment. Statistical significance determined using unpaired student’s t test (**E**). Related to Figure 5

